# Gene knockout studies of Dps protein reveals a novel role for DNA-binding protein in maintaining outer membrane permeability

**DOI:** 10.1101/2024.05.28.596291

**Authors:** Indu Pant, Akhilesh A. Potnis, Ravindranath Shashidhar

## Abstract

DNA-binding proteins like Dps are crucial for bacterial stress physiology. This study investigated the unexpected role of Dps protein in maintaining outer membrane integrity of *Salmonella* Typhimurium. We observed that a *Δdps* mutant displayed increased sensitivity to glycopeptide antibiotics (vancomycin, nisin), which are ineffective against Gram-negative bacteria due to their thick outer membrane (OM). Furthermore, the *Δdps* mutant exhibited susceptibility to membrane-disrupting agents like detergents (deoxycholate, SDS) and phages. The perforation was observed in OM after the treatment of vancomycin using atomic force microscopy (AFM). Notably, this sensitivity was rescued by supplementing the media with calcium and magnesium cations. These findings suggest a novel function for Dps in maintaining outer membrane permeability. We propose two potential mechanisms: 1) Dps might directly localize to the outer membrane, or 2) Dps might regulate genes responsible for lipopolysaccharide (LPS) synthesis or outer membrane proteins, key components of outer membrane. This study highlights a previously unknown role for Dps beyond DNA binding and warrants further investigation into the precise mechanism by which it influences outer membrane integrity in *Salmonella*.

**IMPORTANCE:** Large, hydrophilic glycopeptides like vancomycin are ineffective against Gram-negative bacteria due to their inability to penetrate thick outer membrane of the Gram-negative bacteria. This study investigates the role of the DNA-binding protein, Dps, in maintaining outer membrane integrity. We demonstrate that Dps loss renders bacteria susceptible to vancomycin. These findings suggest Dps as a potential target for developing novel therapeutic strategies, potentially involving combinations with glycopeptide antibiotics, to combat resistant against Gram-negative pathogens.

## INTRODUCTION

*S.* Typhimurium is a foodborne pathogen (1), recognized for its resistance to antibiotics and its ability to endure various environmental conditions (2). The bacterial cell has developed numerous mechanisms to withstand both biotic and abiotic stresses. Among these adaptations, DNA binding proteins play a crucial role in making cells tolerant to stress condition. Various DNA binding proteins (DBPs) have been identified (3) and one such protein is derived from starved cells, known as DNA binding protein from starved cells (Dps). Dps has demonstrated its efficacy in shielding cells against multiple stressors (4). Structurally akin to ferritin, Dps exhibits DNA binding, iron binding, and ferroxidase activities (5). Its expression is regulated by different global gene regulators such as, RpoS, SigmaS, and OxyR (6). Furthermore, the expression of the Dps protein is subject to repression by Fis and HNS proteins (7).

In *Escherichia coli*, upon exposure to oxidative or nutritional stress conditions, the Dps protein is induced, leading to its binding to DNA in a manner lacking apparent sequence specificity. This binding results in the formation of highly stable complexes, thereby safeguarding the DNA against oxidative stress (8). Dps serves a pivotal role in shielding cells from both oxidative and nutrient-related stressors. Numerous studies have delved into the structural and functional characteristics of Dps and homologues of Dps have been identified in over 1000 distantly related bacteria and archaea (9). Hence, the functional role of this protein may be far-reaching.

The Dps protein is primarily expressed during the stationary phase of cellular growth. Bacteria in the stationary phase exhibit reduced susceptibility to antibiotics compared to the exponentially growing cells. Moreover, the stationary phase physiology is associated with transitions to the Viable But Nonculturable (VBNC) and persistence states under conditions of nutrient starvation (10). Therefore, we thought proteins like Dps which are present in abundance in stationary might have a role in deciding cell’s sensitivity to certain class of antibiotics. Hence, in this investigation, we aimed to assess alterations in sensitivity towards antibiotic classes that are typically ineffective against Gram-negative bacteria. Our focus was on vancomycin, a glycopeptide antibiotic often utilized in the treatment of Gram-positive bacteria. The thick outer membrane of Gram-negative bacteria renders them impermeable to high molecular-weight antibiotics like vancomycin (11). Despite efforts to modify vancomycin for efficacy against Gram-negative bacteria, such attempts have met limited success. Vancomycin exerts its antimicrobial activity by inhibiting cell wall synthesis through binding to the D-alanyl-D-alanine (D-Ala-D-Ala) peptide motif (12).

In the current study, we found that removal of Dps protein made Δ*dps* mutant sensitive to vancomycin antibiotic, hence Dps protein might be important for maintaining outer membrane permeability. Hence, we further delve into understanding the underlying mechanisms responsible for the vancomycin sensitivity of Dps null mutants.

## RESULTS

### Confirmation of *Δdps* deletion strain

The Δ*dps* strain was successfully constructed. The transformants were selected by plating on LB agar plates containing 50µg/ml of kanamycin antibiotic. The correct insertion of kanamycin cassette in the place of *dps* gene was confirmed using PCR. The whole genome sequencing of knockout strain confirmed the in-frame deletion of *dps* gene. The complemented strain was also confirmed by PCR and restriction digestion. Further, all the phenotypes observed in the mutant with respect to wild type were restored in the complemented strain.

### Sensitivity of *Δdps* to vancomycin

The stationary phase bacteria are less susceptible to antibiotics than their exponentially growing counter parts. Therefore, the idea was to check for change in sensitivity towards antibiotics after removal of *dps* gene. The MIC of vancomycin for *S*. Typhimurium was determined using microbroth dilution assay (Fig. 1a). The elimination of *dps* gene from the wildtype strain lead to its increase in sensitivity to vancomycin (Fig. 1b). No growth was observed in LB medium substituted with vancomycin after overnight incubation (Fig. 1c). It was observed that in the presence of 1mg/ml vancomycin in the medium, the cells of Δ*dps* were getting killed, whereas for wild type *S*. Typhimurium MIC was >1mg/ml. The vancomycin was having bactericidal effect on the mutant as no mutant was recovered after plating on Luria agar plates.

**FIG 1.**
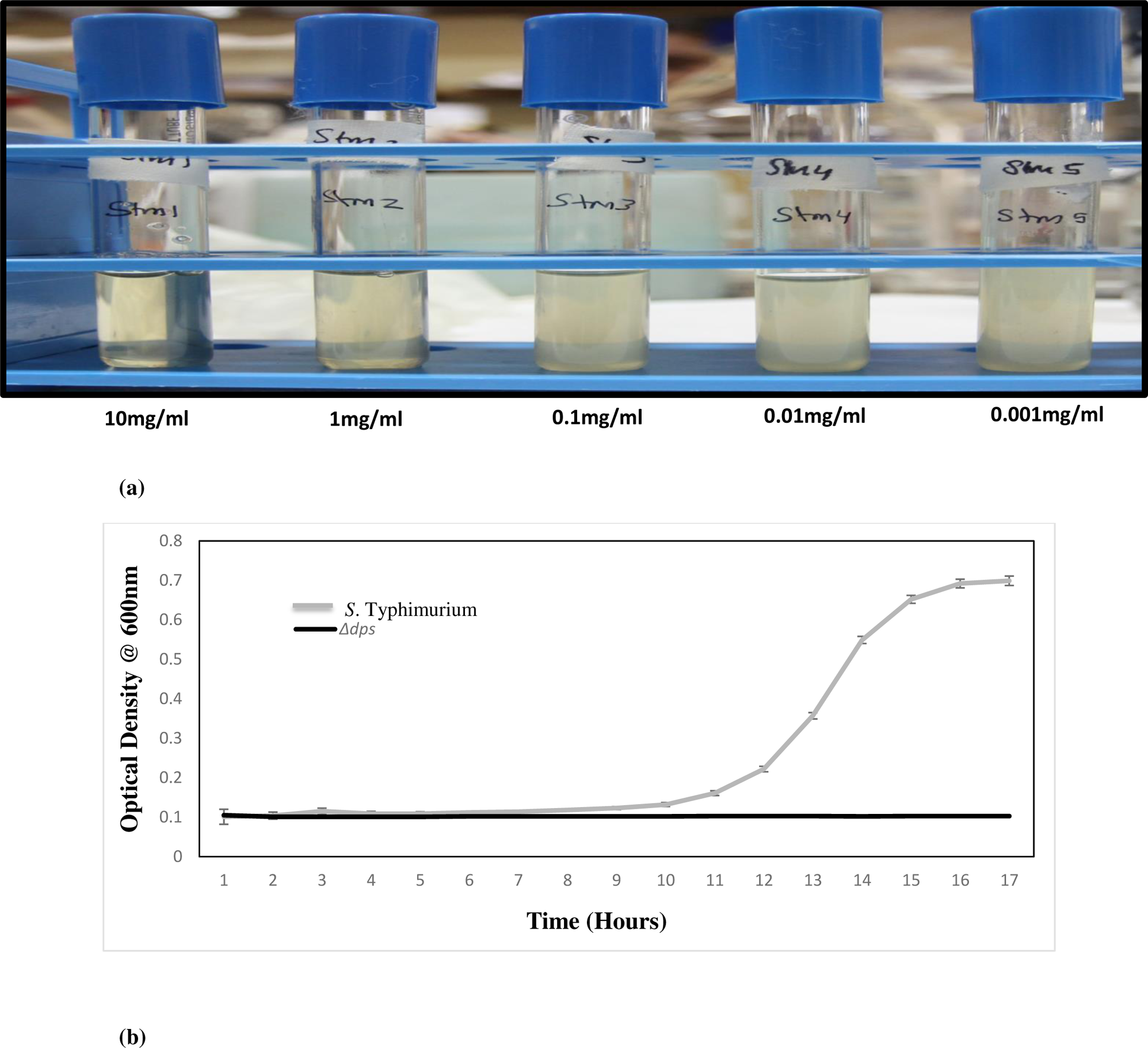
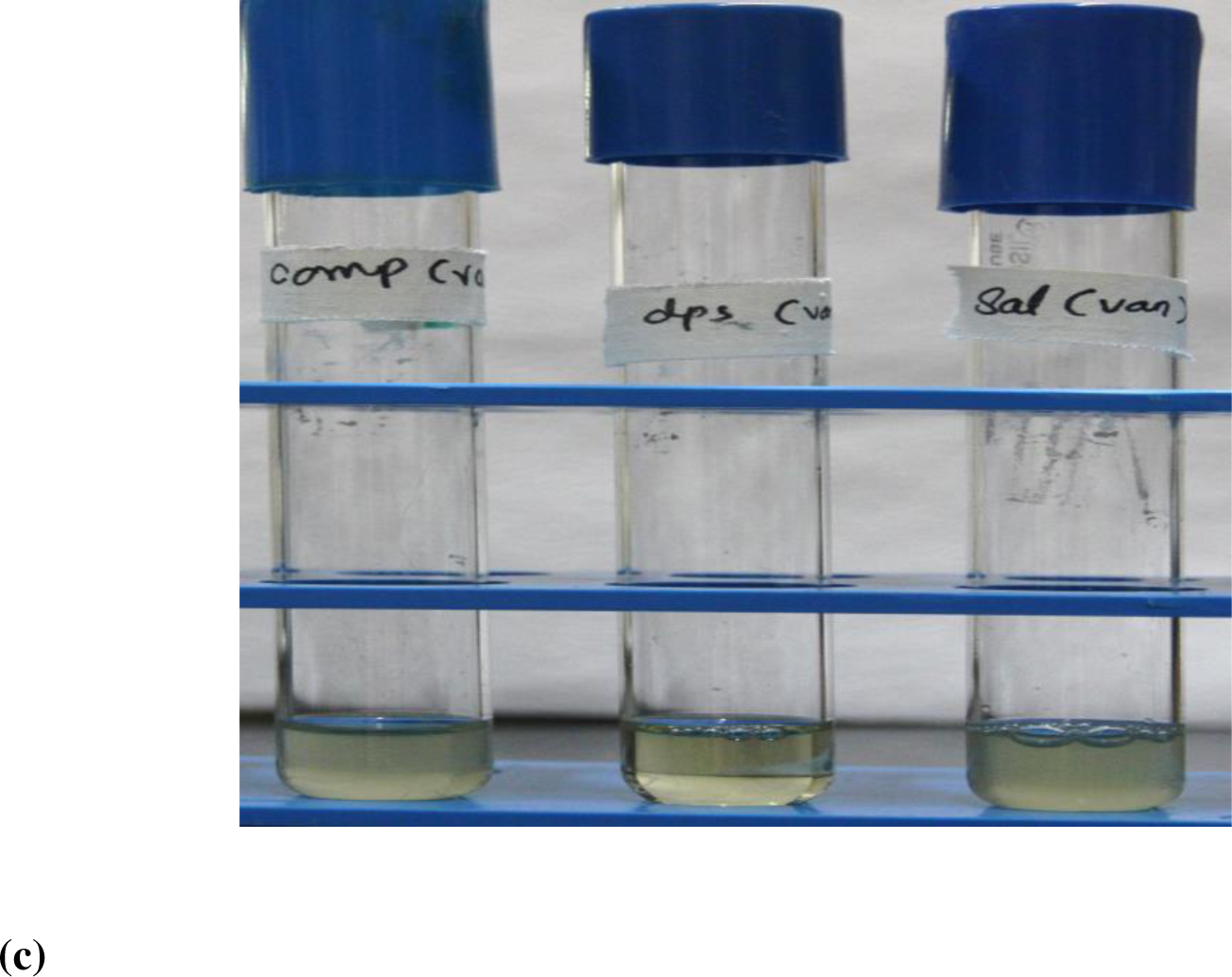
(a) The MIC of vancomycin for *S*. Typhimurium was determined using microbroth dilution assay. (b) The effect of vancomycin on the growth of *S.* Typhimurium and *Δdps* was observed by growing wild type and mutant strain in the Mueller Hinton medium in the presence of vancomycin antibiotic (1mg/ml). The *Δdps* did not grow in the medium in the presence of antibiotic. (c) No growth observed in vancomycin substituted medium after overnight growth in *Δdps;* whereas wild type *Salmonella* and *dps\** grew well.

### Sensitivity of *Δdps* to other antibacterial agents

Since the mutant displayed sensitivity to a drug typically effective against Gram-positive pathogens, we aimed to assess its sensitivity to similar drugs. Consequently, we selected nisin for this purpose. Both the mutants and wild-type cells were exposed to nisin at a concentration of 100µg/ml. Remarkably, our findings revealed that in the presence of nisin, the *Δdps* mutant perished, whereas the *S*. Typhimurium strain remained unaffected (Fig. 2).

**FIG 2.**
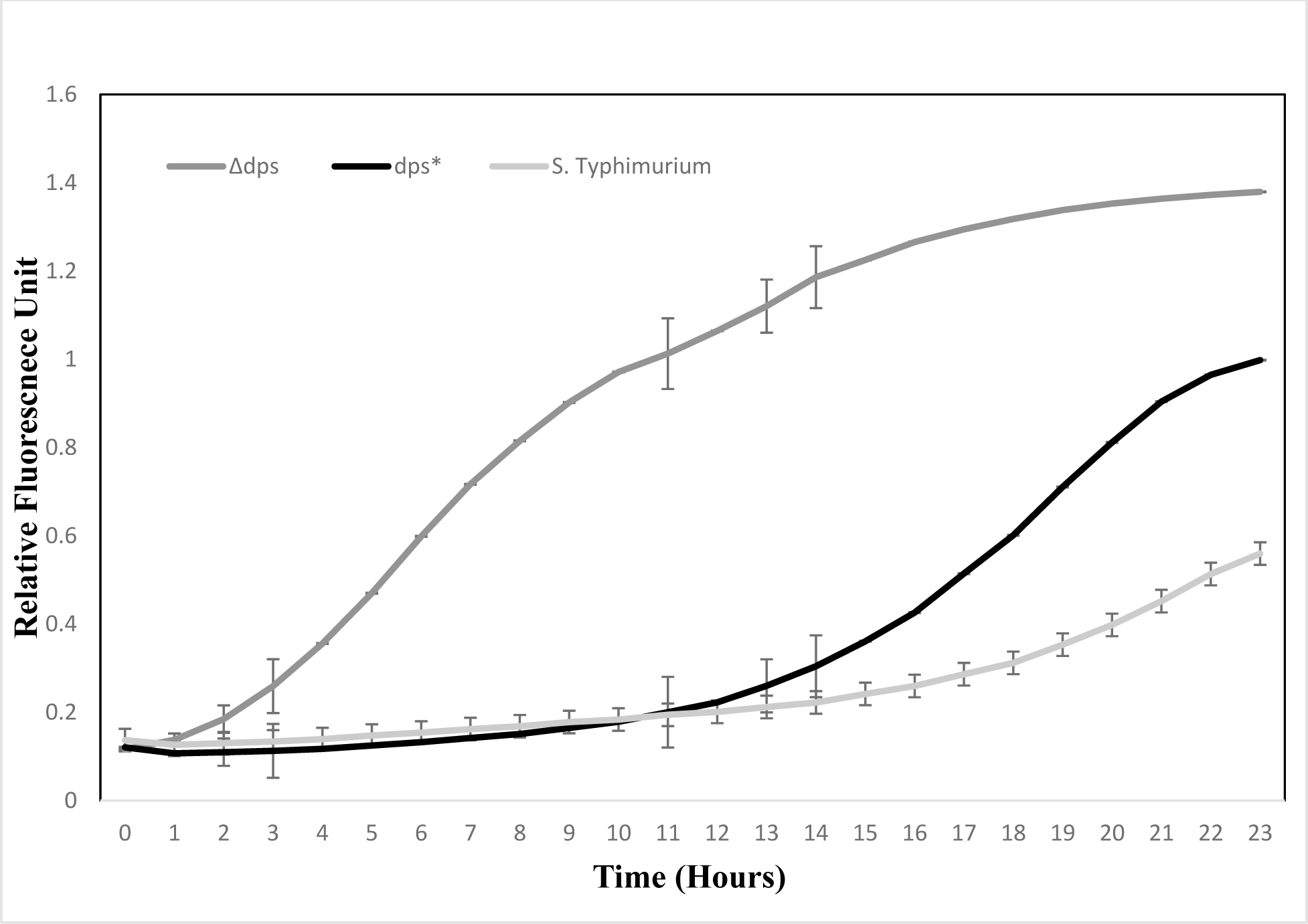
The wild type *S*. Typhimurium, *Δdps* and *dps\** were treated with nisin (100µg/ml) and relative fluorescence unit were measured using uptake of propidium iodide dye. *Δdps* was found to be most sensitive to the nisin treatment

### Sensitivity of *Δdps* to membrane disrupting compounds

The effect of different membrane disrupting compounds were also observed on the Δ*dps* strain (13). The Δ*dps* strain were showing sensitivity to 0.5% deoxycholic acid (Fig. 3b) and 2% SDS (Fig. 3c).

**FIG 3.**
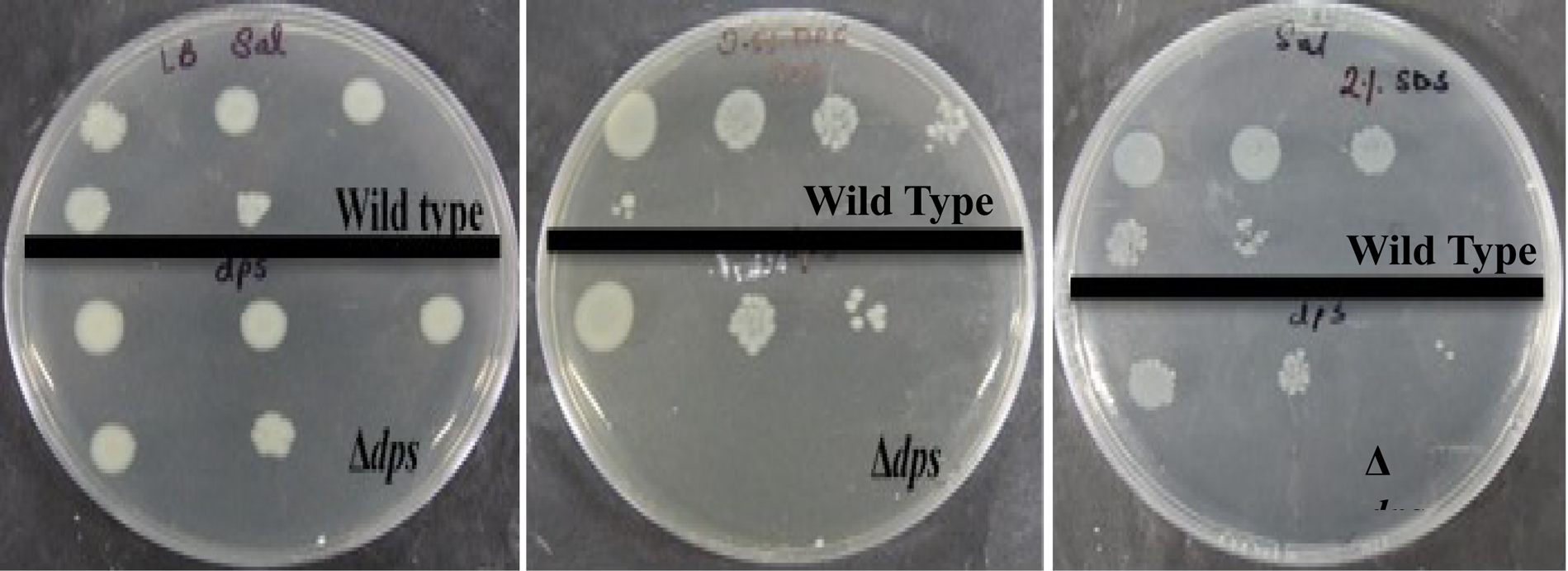
Effect of membrane disrupting compound on *S.* Typhimurium and Δ*dps.* The upper panel represent wild type *S*. Typhimurium and lower panel represent mutant Δ*dps* (a) The wild-type and Δ*dps* mutant cells were serially diluted from 10^8^ to 10^3^ cells/ml in 10-fold steps and spotted onto a LB plate. (b) The wild-type and Δ*dps* mutant cells were serially diluted from 10^8^ to 10^3^ cells/ml in 10-fold steps and spotted onto a LB plate supplemented with 0.5% deoxycholic acid (c) The wild-type and Δ*dps* mutant cells were serially diluted from 10^8^ to 10^3^ cells/ml in 10-fold steps and spotted onto a LB plate supplemented with 2% SDS.

### Sensitivity of *Δdps* to phages

We investigated the sensitivity of the *Δdps* strain towards phages (14). Notably, our results revealed that the Δ*dps* strain exhibited heightened sensitivity to both PM9 and PM10 phages (Fig. 4a, 4b).

**FIG 4.**
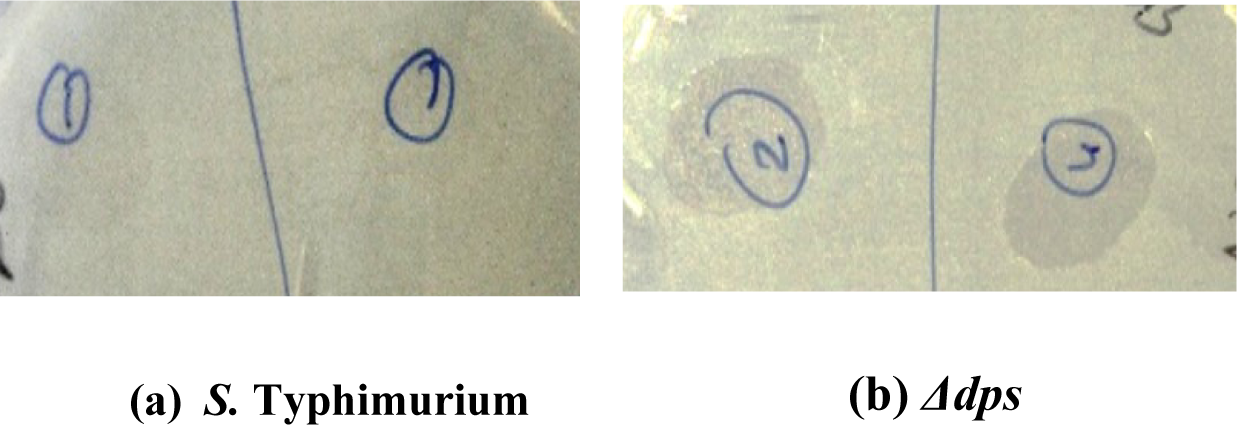
The bacterial lawn was allowed to grow on LB agar plates. 10ul spot of Phage PM9 and PM10 was put on the bacterial lawn (a) *S.* Typhimurium (b) *Δdps* to check for sensitivity towards phages. *Δdps* was very sensitive as evident from zone of clearance made, whereas no such zone was observed in wild type *Salmonella*.

### Restoration of antibiotic sensitivity in *Δdps*

The bridge between a protein and the cell membrane is formed by calcium ions (15). Calcium ions are known for stiffening and maintaining the arrangement of lipid bilayer (16). Therefore, the rigidity of outer membrane can be changed in *S.* Typhimurium and *Δdps* using Ca^+2^ and Mg^+2^. The parent and mutant was exposed with 10 mM and 20 mM calcium chloride and magnesium sulphate and it was found that mutants treated with 10 mM and 20 mM calcium chloride (Fig. 5a,5b,5c), and magnesium sulphate (Fig. 6a,6b,6c), were showing resistance to vancomycin, thus restoring the phenotype similar to that of wild type *S*. Typhimurium.

**FIG 5.**
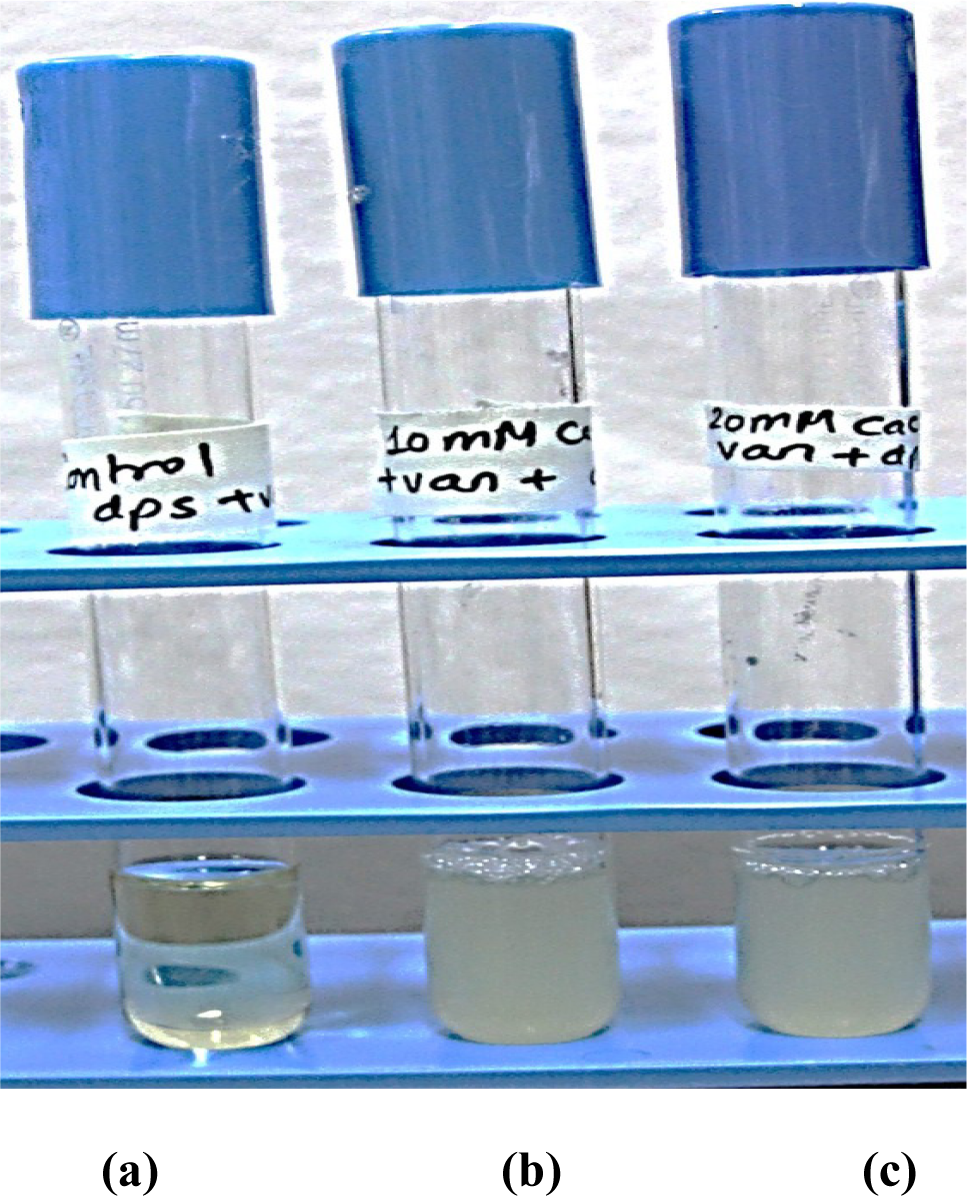
The mutant was treated with cations like calcium and then exposed to vancomycin antibiotic. (a) *Δdps* +vancomycin; (b) *Δdps* +vancomycin+10mM Cacl_2_; (c) *Δdps* +vancomycin+20mM Cacl_2_.The *Δdps* mutant became resistant to vancomycin antibiotic after treatment with calcium chloride, as Calcium chloride is known to stabilize and maintain integrity of outer membrane

**FIG 6.**
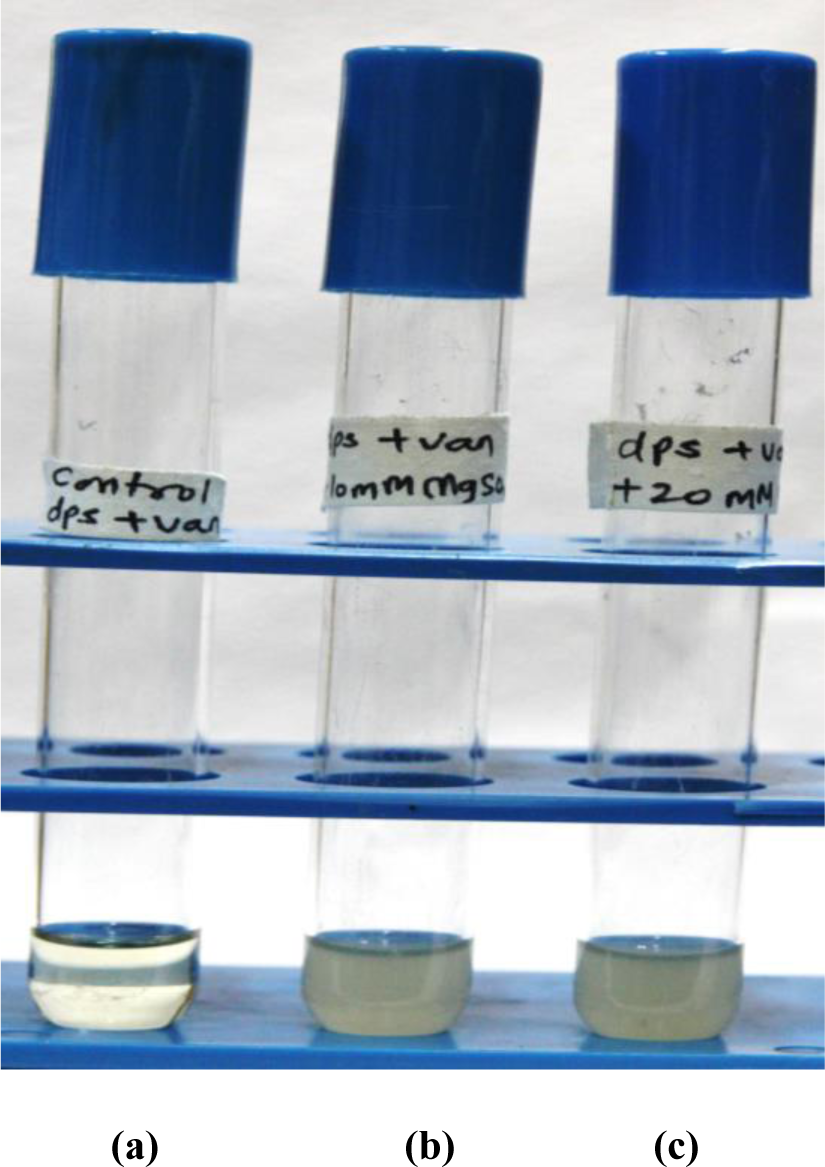
The mutant was treated with cations like magnesium and then exposed to vancomycin antibiotic. (a) *Δdps* +vancomycin; (b) *Δdps* +vancomycin+10mM MgSO_4_;(c)*Δdps* +vancomycin+20mM MgSO_4_.The *Δdps* mutant became resistant to vancomycin antibiotic after treatment with calcium chloride, as Calcium chloride is known to stabilize and maintain integrity of outer membrane

The amount of Mg^+2^ and Ca^+2^ was estimated in both wild type and mutant strain and it was found that there was non-significant difference in the concentration of both the ions. In wild type, Mg^+2^ was 1.36 ppm and in Δ*dps* was 1.53 ppm (Table 3). Similarly, in both wild type and Δ*dps,* Ca^+2^ was <0.5ppm.

### Sensitivity of wild type to vancomycin in presence of chelators

We found that *Δdps* was sensitive to different antimicrobials and detergents and we anticipated that the reason behind the enhanced sensitivity was changes in OM permeability. Therefore, to further validate our hypothesis, we tried to change the OM permeability in wild type strain by treating wild type cells with chelators like EDTA, that destabilizes the outer membrane by removing cations like Mg^+2^ and Ca^+2^. *S*. Typhimurium was found to be sensitive to vancomycin in the presence of EDTA (Table 1); which showed that changing OM permeability indeed changes sensitivity of wild type to vancomycin.

**Table 1.**
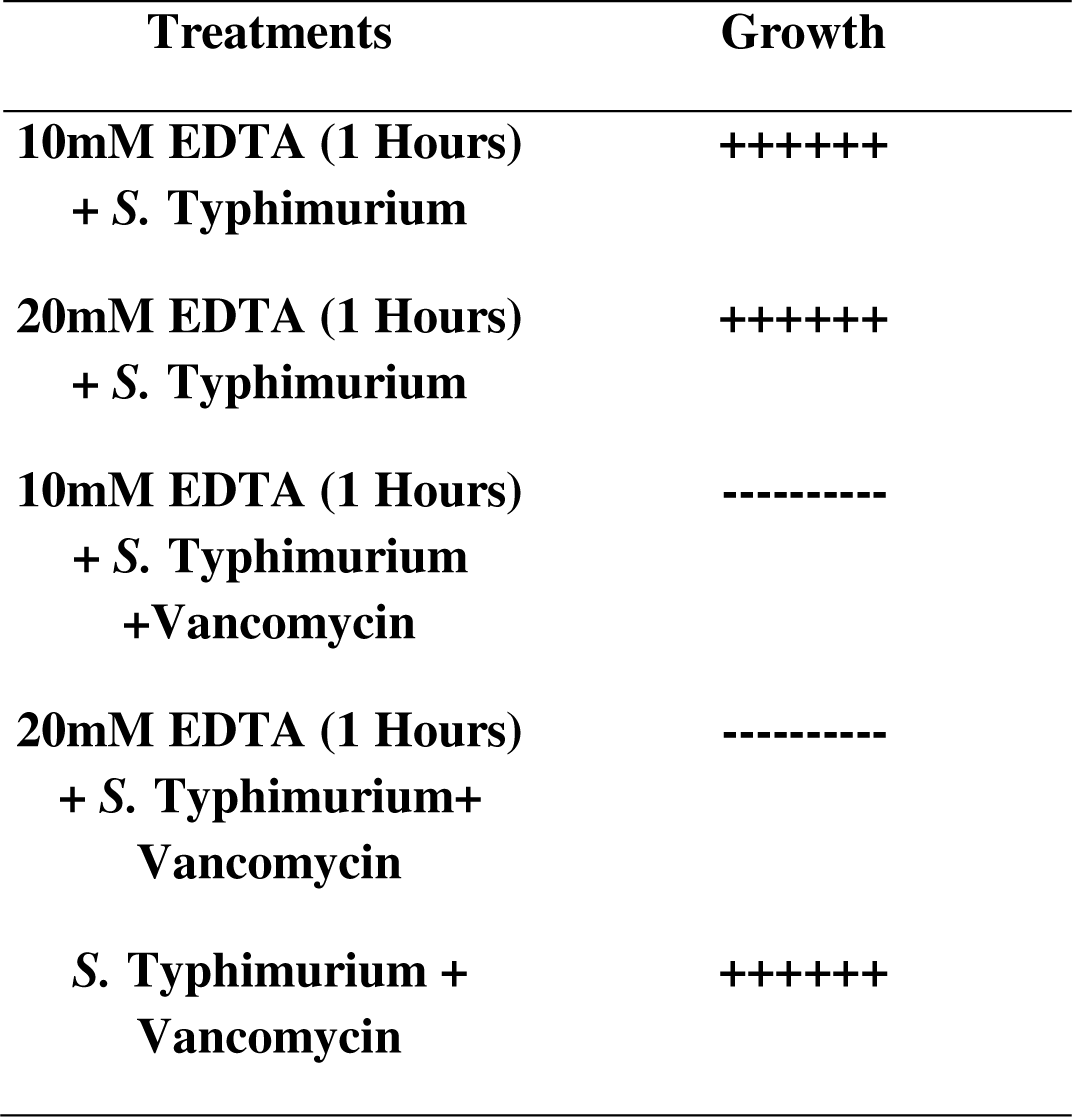
The table show growth of *S.* Typhimurium as observed after different treatments. -- represent no growth; ++++++ represent confluent growth. This showed that in presence of EDTA cells were able to grow, so EDTA at 10mM and 20mM is not toxic to cells but on addition of vancomycin in presence of EDTA cells growth did not happen as cells outer membrane integrity was changed leading to accumulation of big molecule like antibiotic inside the cells and hence death of *Salmonella* which result in no growth.

| **Treatments** | **Growth** |
| --- | --- |
| **10mM EDTA (1 Hours)**<br>**+ *S. Typhimurium*** | **++++++** |
| **20mM EDTA (1 Hours)**<br>**+ *S. Typhimurium*** | **++++++** |
| **10mM EDTA (1 Hours)**<br>**+ *S. Typhimurium***<br>**+Vancomycin** | **-----** |
| **20mM EDTA (1 Hours)**<br>**+ *S. Typhimurium*+**<br>**Vancomycin** | **-----** |
| ***S. Typhimurium* +**<br>**Vancomycin** | **++++++** |

### Sensitivity of wild type to vancomycin at higher temperature

It was observed that wild type became sensitive to vancomycin at 42℃ (Table 2). This might have happened as change in temperature leads to changes in outer membrane integrity leading to wild type *S*. Typhimurium showing sensitivity to vancomycin (17).

**Table 2.** Treatment of *S.* Typhimurium and *Δdps mutant* with vancomycin at higher temperature showed that no growth of wild type at higher temperature occurs in presence of vancomycin which is the result of change in membrane integrity.

| Treatment with vancomycin at<br>42°C | Occurrence of Growth |
| --- | --- |
| *S. Typhimurium*+ Vancomycin | ---- |
| *Δdps* + *Vancomycin* | ---- |

### The integrity of outer membrane is compromised

As depicted from all the above-mentioned results, membrane integrity is compromised in *Δdps*.

We tried to detect dissipation of the transmembrane potential in Δ*dps*; by using membrane depolarization dye like DiSC_3_. It was found that complete depolarization of the outer membrane in the presence of vancomycin occurred in Δ*dps,* whereas no such depolarization was observed in the case of wild type *S.* Typhimurium (Fig. 7). DiSC_3_ is a cationic dye and it gets incorporated within the hydrophobic membrane but once vancomycin is encountered by the cell, membrane disruption will occur leading to release of DiSC_3_ in the medium hence increase in the relative fluorescence unit.

**FIG 7.**
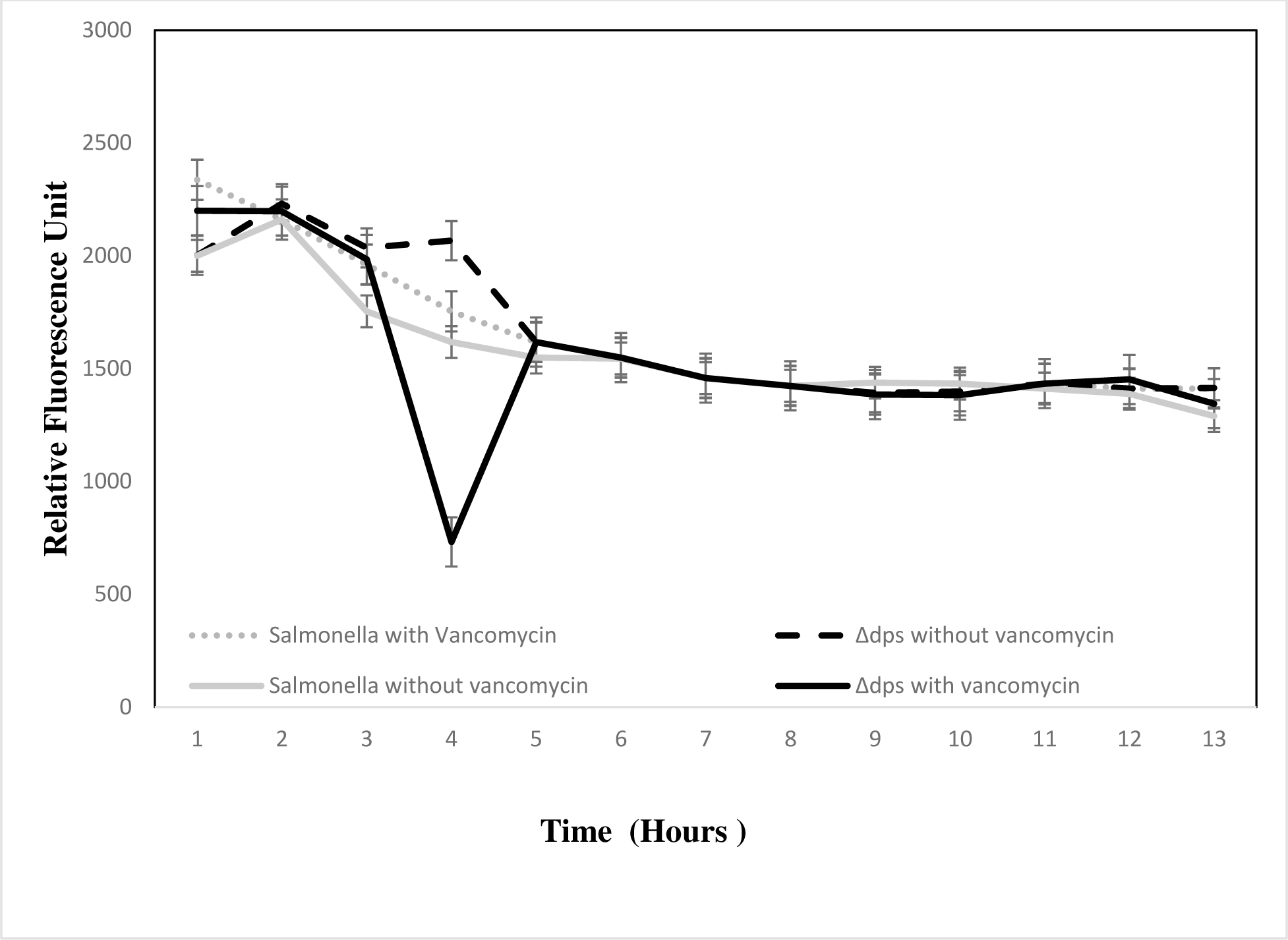
Membrane potential sensitive fluorescent dye DiSC_3_. Fluorescence intensity changes of DiSC_3_ in a cell suspension of *S*. Typhimurium and Δ*dps* on addition of vancomycin antibiotic (1mg/ml). Fluorescence intensity changes of DiSC 3 in a cell suspension of *S*. Typhimurium and Δ*dps* without the addition of vancomycin antibiotic.

**Table 3.**
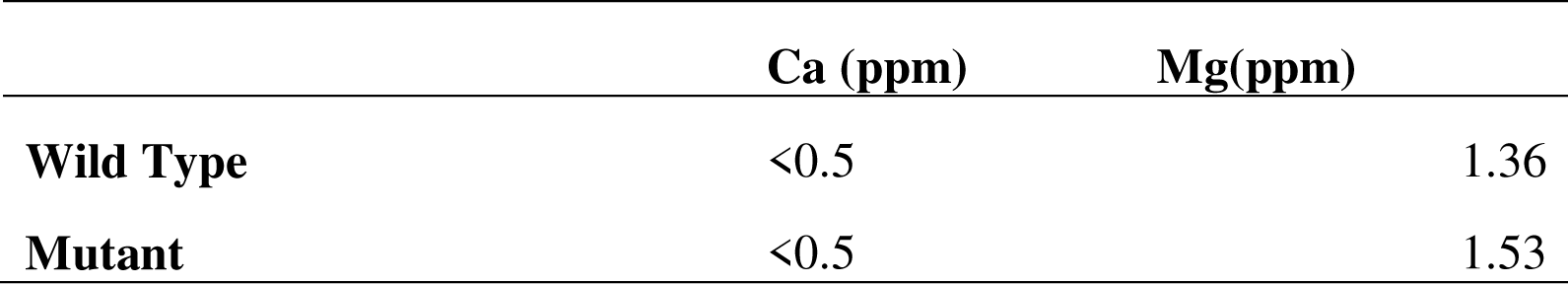
Estimation of calcium and magnesium ions in wild type *S.* Typhimurium and *Δdps* using ICP-MS.

To reveal the changes in dynamic structure of microbial membrane integrity after treatment with vancomycin, Atomic Force Microscopy was also employed and images showed the presence of pores on the surface of Δ*dps* after vancomycin treatment whereas no such pores were found on the surface of wild type *S*. Typhimurium (Fig. 8a-c). The formation of pores on Δ*dps* (Fig. 8 d) clearly indicated the cells entering into lysis state.

**FIG 8.**
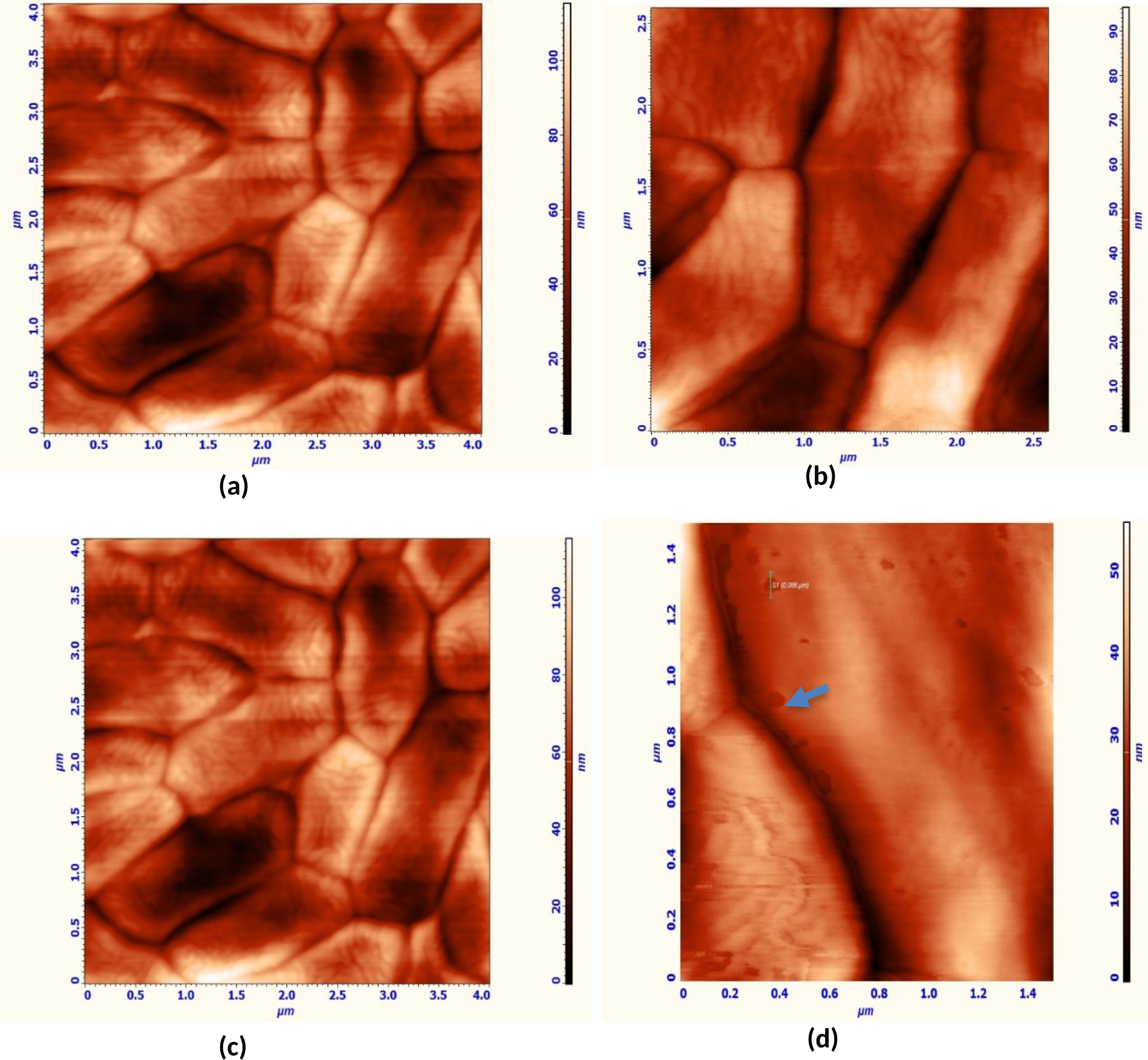
AFM analysis was performed to observe changes in the membrane on the application of vancomycin: (a) Wild type *S*. Typhimurium cells; (b) Wild type *S*. Typhimurium cells after vancomycin treatment); (c) Δ*dps* without vancomycin treatment; (d) Δ*dps* with vancomycin treatment (1mg/ml), the pore formed by vancomycin treatment is marked by arrow.

### Gene expression studies to identify the role of *dps* gene in enhancing sensitivity of S. Typhimurium to vancomycin

The transcriptome analysis was carried out between wild type *S*. Typhimurium and Δ*dps* mutant in Luria Broth medium in exponential phase. The results are presented in Table S2; it represents some of the important genes which were highly upregulated in transcriptome analysis. The comparison of the mean levels of expression of sample duplicates between wild type and mutant was used for evaluating differential expression of genes. At P value of <0.01, differentially regulated genes were studied if fold change was >1.5 or <-1.5.

The major groups to which differentially regulated genes were clustered belonged to metabolism, membrane proteins, DNA repair and pathogenesis.STM4217 (Lytic murein transglycosylase), STM3026 (Outer membrane protein), STM2102 (Lipopolysaccharide biosynthesis protein WzxC), PSLT017 (Outer membrane usher protein PefC), STM2099 (colanic acid biosynthesis protein,WcaM), STM4097 (outer membrane protein), STM1131 (Outer membrane protein), STM1043 (Outer membrane protein X),STM 2514 (Outer membrane protein) were upregulated in Δ*dps.* The gene expression analysis (Real time PCR) showed an upregulation of many membrane-associated proteins STM2102, STM1043, PSLT017, STM2756, STM2151, STM2514, STM3026, STM2102 and STM1131 which is in sync with the transcriptome data (Fig. 9). In most of the cases, there were 4-fold upregulation of the genes.

**FIG 9.**
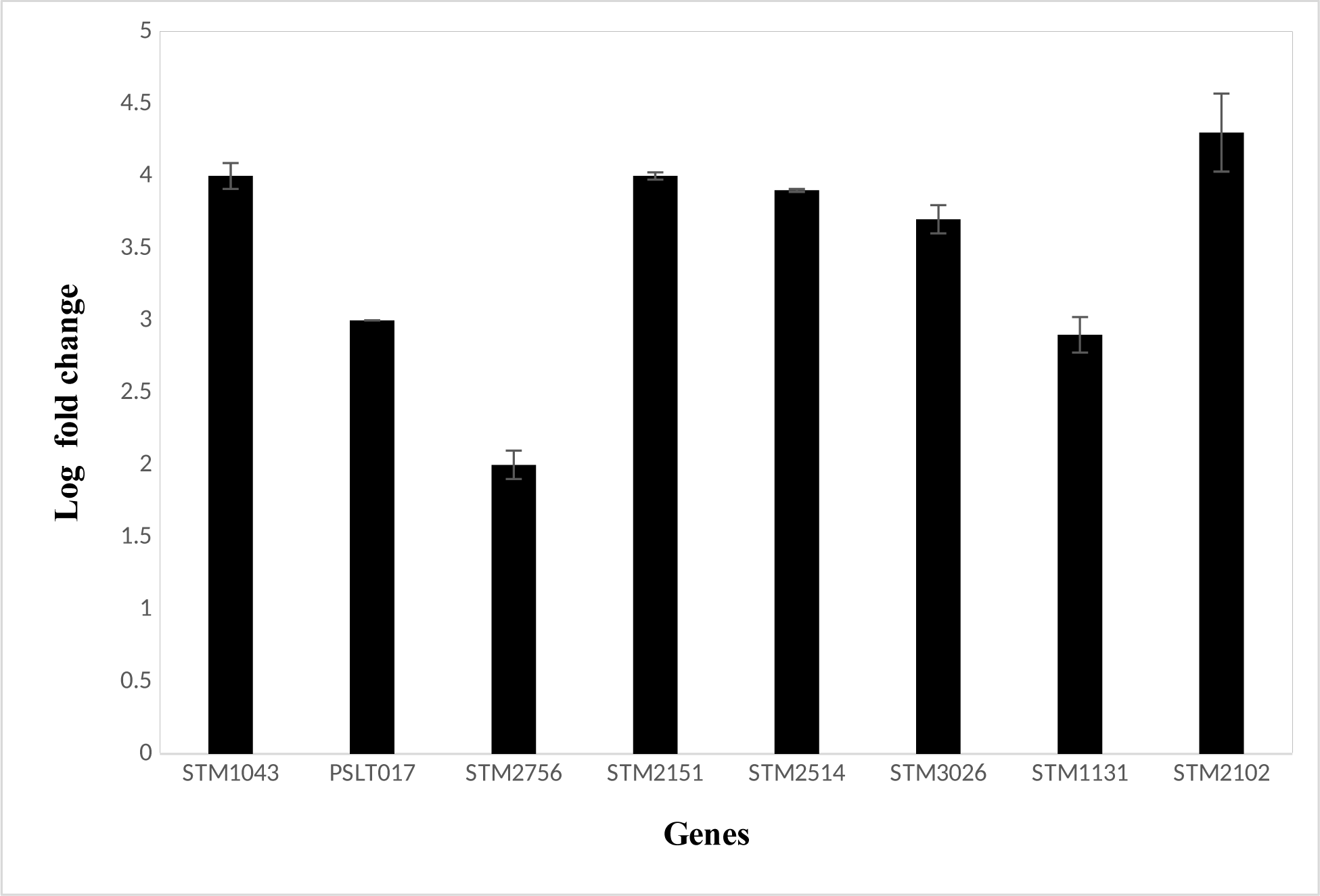
The cells of wild type S. Typhimurium and mutant were allowed to grow till exponential phase and RNA was isolated using TRIzol reagent; after cDNA synthesis, RT-PCR analysis of different genes associated with outer membrane and LPS were performed. The graph is representing log fold changes in the expression of genes.

## Discussion

Multiple molecular and physiological functions are attributed to nonspecific DNA binding proteins. In this study, our focus was to investigate the change in outer membrane permeability upon the successful knockout of the *dps* gene; as gram negative pathogens are resistance to vancomycin.

In our experiments, we observed that the elimination of the *dps* gene from the wild-type *S.* Typhimurium genome increased its sensitivity to vancomycin. We found that *Δdps* cells exhibited a 10-fold higher sensitivity to vancomycin than the wild type (Fig. 1). The vancomycin is a large glycopeptide antibiotic. Gram-negative bacteria are typically resistant to this antibiotic due to the impermeability of their thick outer membrane (11). However, the *Δdps* mutant displayed sensitivity to vancomycin, suggesting a potential change in the integrity of its outer membrane, enabling the antibiotic to act on the peptidoglycan in this knockout strain. The peptidoglycan layer can only be accessible to large hydrophobic molecules like vancomycin after some defects in outer membrane.

To confirm whether the increased sensitivity was indeed due to outer membrane damage, we conducted further investigations. We took another antibacterial agent which is similar to vancomycin in mode of action against Gram negative pathogens. The antibacterial agent was nisin. We observed that the mutant was sensitive to nisin. This experiment proved that due to change in outer membrane structure in Gram negative bacteria is responsible for sensitivity to nisin and similar observation regarding outer membrane has been reported earlier (18).

The sensitivity of mutant and wild type was also evaluated against other OM disrupting compounds like 2% SDS, 0.5% deoxycholic acid and phages. The mutant was more sensitive to these compounds and phages which again point towards change in membrane permeability in Gram negative bacteria. The changes in permeability has led to more damage in mutant by detergents and phages to which wild type *S.* Typhimurium was earlier not susceptible. The Gram-negative bacteria are generally resistant to action of detergents because of their unique cell wall structure. The outer leaflet of outer membrane of Gram-negative bacteria contain lipopolysaccharide which help cells to abstain passive diffusion of hydrophobic molecules like detergents (19). So, if gram negative cells are becoming sensitive to detergents, it depicts changes in barrier provided by LPS. It had already been shown in *E. coli,* that wild type strain and mutants have different susceptibilities to agents like deoxycholate and mutant become susceptible to T-even phages due to alterations in the integrity of their outer membrane (20).

This sensitivity to vancomycin could be replicated by treating wild type with cation chelating agent like EDTA (17). The EDTA removes cations from outer membrane, which would impair the integrity of OM. Similarly, supplementing medium with cations like calcium and magnesium rescued the sensitivity of *Δdps* towards vancomycin. The divalent cations are capable of binding to adjacent LPS molecules and after binding to negatively charged lipid molecule, it neutralizes the repulsive forces and helps LPS molecules stay together (21) and hence stabilizes the outer membrane but chelators like EDTA disrupt the arrangement of LPS molecules.

The OM integrity also could be modified by changing the incubation temperature (22). At higher temperature unsaturation level in the membrane lipid decreases (23), this lead to changes in membrane integrity. In 2016, it had been shown that temperature alter the structure of membrane lipid (24). When wild type cells were incubated at 42℃ and treated with vancomycin, the wild type cells become as sensitive as *Δdps* mutant to vancomycin. Increase in temperature changes sensitivity of *Staphylococcus* towards antibiotics like daptomycin, vancomycin, tigecycline, fosfomycin, and cefamandole (25,26). These antibiotics belong to different classes, which mean irrespective of the class of antibiotics, temperature increases sensitivity towards antibiotics. This suggests that, increase in temperature changes the outer membrane fluidity. Thereby, the antibiotics could go through outer membrane. The fluorescence assay using the membrane potential-sensitive probe DiSC_3_ depicted vancomycin treatment induced membrane potential dissipation in the *Δdps* strain, but not in wild-type *S.* Typhimurium, which again point towards changes in membrane permeability.

The transcriptome and real time PCR analysis also showed upregulation of outer membrane proteins and lipopolysaccharide synthesis genes which signifies changes in membrane component in *Δdps* strain, giving another hint in change in membrane permeability in the mutant strain.

Can the outer membrane permeability change lead to damage? All the above experiments were physiological proof for the damage. However, that is not the direct evidence. Hence, we analyzed the cells by AFM. Our results were similar to results obtained by Meincken et al. (27). In our study, the vancomycin treated cells showed significant perforation on the surface. This gives clear evidence that indeed the outer membrane damage is there in the presence of vancomycin in *Δdps*.

How does DPS protein is linked to the change in outer membrane integrity? Is it the result of a DNA binding activity or is it a result of any other gene mutation happened during the knockout experiment? To rule out this possibility we carried out two experiments. The first is the complementation. The mutant was complimented with wild type *dps* gene with the native promoter in Pbad-Topo vector. This complementation rescued the antibiotics sensitivity. The whole genome sequence of the knockout was done to identify any other mutation (data not shown) that might be giving this phenotype and we did not find any nonsynonymous mutations.

To link the DPS protein to outer membrane integrity, we considered two possibilities. The first is that DPS itself gets integrated into the outer membrane, as some studies have reported the presence of a fraction of DPS protein in the outer membrane (28, 29).Therefore, supplementing the membrane with the metal ions like Ca^+2^ and Mg^+2^ restored vancomycin resistance, supporting the possible presence of DPS in outer membrane and in absence of *dps*, stabilization by the presence of cations.

The second possibility is that, DPS involved in regulation of outer membrane genes or genes involved in the synthesis of LPS, because outer membrane proteins and LPS are associated with antibiotic resistance phenotype. The transcriptome data did give some clues (Table S2). We found that 8 of the outer membrane genes were upregulated in the mutant. The up or down regulation of the outer membrane gene would lead to destabilization of the membrane.

Further experiments are needed to unravel entire mechanism of Dps protein in maintaining outer membrane stability. Our results suggest moonlighting effect of this DNA binding protein.

### Implication

Current study investigated the role of a DPS in protecting *S.* Typhimurium from the antibiotic vancomycin. We found that removing the *dps* gene made the bacteria more sensitive to vancomycin, suggesting a weekend outer membrane integrity. Our experiments suggested that *dps* may be influencing the expression of certain genes involved in membrane structure or it may be itself present in the outer membrane. Hence, *dps* plays a part in maintaining the integrity of the outer membrane in Gram negative bacteria like *S*. Typhimurium.

## MATERIALS AND METHODS

### Bacterial strains, reagents and growth conditions

The knockout of *dps* was constructed in wild type strain of *Salmonella* enterica subsp. enterica serovar Typhimurium strain LT2 (MTCC 98) procured from Microbial Type Culture Collection and Gene Bank (MTCC) Chandigarh, India. The growth-related experiments were carried out in Lysogeny broth (LB) (HiMedia, India). LB agar plates supplemented with kanamycin (50 µg/ml) or carbenicillin (50 µg/ml), vancomycin (1mg/ml) were used as per requirement. All the reagents were autoclaved or filter sterilized before use.

### Construction of gene knockout mutant and complementation of *Δdps* strain

The construction of *dps* genes mutant in *S*. Typhimurium was done by the Quick and Easy *E. coli* Deletion Kit by Red®/ET® Recombination, (Gene Bridge, Heidelberg, Germany). The sequence for *dps* gene was taken from NCBI (NC_003197.2). The LB plates supplemented with kanamycin antibiotic (50µg/ml) was used for selection of knockouts. The knockouts were confirmed using PCR based methods and whole genome sequencing (Genotyping Tech. Pvt. Ltd.). The phenotypic characteristic of mutant was compared with the phenotype given in literature. The pBAD TOPO™ TA Expression Kit, Invitrogen, Massachusetts, United States was used for constructing *dps* gene complement strain with the native promoter (200 bp upstream region to the *dps*) in the Δ*dps* mutant. The *dps* complemented strain is referred to as *dps*\* in the subsequent section. The primers and their sequences used in the study are given in Supplementary data Table S1.

### Vancomycin sensitivity assay

The wild type *S.* Typhimurium and knockout strain of Δ*dps* were grown overnight. From overnight grown culture, 1:100 cells were re-inoculated into fresh LB medium till exponential phase and 10^5^ cells/ml of wild type and Δ*dps* were exposed to vancomycin (1mg/ml) in 5ml of Mueller Hinton broth (MHB) medium. The tubes were incubated at 37℃ overnight. Tubes were observed for presence or absence of turbidity the next day.

### Effect of nisin on sensitivity of *Δdps*

Overnight cells of *S.* Typhimurium, Δ*dps* strains were taken. These were washed with saline and then resuspended in phosphate buffer saline (PBS) having 100 µg/ml nisin and death of the strains in the presence of nisin were measured with the uptake of propidium iodide. The relative fluorescence unit was measured in Bio-Tek Synergy H1 plate reader (Winooski, Germany) for a period of 16 hours.

### Effect of EDTA, calcium chloride and magnesium sulphate on *S*. Typhimurium’s sensitivity to vancomycin

The wild type *S.* Typhimurium, Δ*dps* was grown overnight. From overnight grown culture, cells were reinoculated into fresh LB medium till exponential phase. The wild type *S*. Typhimurium cells were then exposed with different concentration of EDTA (10 mM, 20 mM) for 1 hour and Δ*dps* were then exposed to Cacl_2_ and Mgcl_2_ (10 mM and 20mM) and then exposed to 1mg/ml vancomycin.

Estimation of calcium and magnesium ions in *S*. Typhimurium and *Δdps*

The cells of *S*. Typhimurium and *Δdps* were grown overnight in Luria broth medium. For magnesium and calcium determination, ICP-MS method was used (30).

### Membrane depolarization studies using DiSC_3_

Overnight grown cells of *S*. Typhimurium and Δ*dps* strains were taken. The membrane depolarization experiments were performed in presence of 1µM DiSC_3_ as described previously (31).

### Phage sensitivity assay

The overnight culture of wild type and Δ*dps* were taken. These were spread plated on LB plates and then phages PM9 and PM10 were used for spot test analysis (14). The plates were incubated at 37℃, overnight. The plates were observed next day for formation of zone of clearance.

### Effect of membrane disruption compounds on mutants

The wild type *S*. Typhimurium and Δ*dps* were grown in LB for 16-18 hours. The stationary phase cells grown in LB were serially diluted from 10^8^ to 10^4^ cells/ml. For spotting on the agar plate, 2 μl of the culture was put onto LB plates supplemented with 2% SDS, 0.5% sodium deoxycholate. The results were analyzed after 16 hours of incubation at 37°C and data stored as photographs taken with a digital camera (13).

### AFM analysis

Sample preparation, measurements and analysis for carrying out Atomic Force Microscope (AFM) studies were performed as described earlier (32).

### Gene expression analysis through Transcriptomics and RT PCR

The wild type *S*. Typhimurium and *Δdps* were inoculated in LB medium and allowed to grow till their number reach 10^9^ cells. 1:100 cells were then reinoculated into fresh LB medium and allowed to harvest after a period of 2 hours (log phase, OD 0.4). The samples for transcriptomics and RT PCR were made as described previously (33,34).

### Statistical Method

All the experiments were performed in triplicates and all the experiments were repeated three times. Standard deviation (SD) was used to analyze real-time PCR results, performed in Microsoft Excel. Significant results were obtained at P<0.05 by performing two sample t-test analysis.

## SUPPLEMENTAL MATERIAL

Table S1 and S2 is available as supplementary material.

## DATA AVAILIBILITY

Transcriptome and whole genome sequencing data will be available on request.

## ACKNOWLEDGEMENTS

Authors want to convey their kind regards to Department of Atomic Energy for their support for providing all the facilities necessary during this work.

